# Utilization of network graph connectivity to evaluate complications of inpatient medical and surgical populations as a method to prioritize quality improvement efforts

**DOI:** 10.1101/099150

**Authors:** Kattie Bear-Pfaffendorf

## Abstract

Network graphs can provide a quantitative framework for identifying complications with significant volumes and strong relationships to other complications, as a method for prioritization of quality improvement work. Here we examine the application of network graphing techniques to acute care inpatient complications on acute care medical-surgical units of a quaternary care center. The 3M PPC software identified 66 complications among 106 unique patients with two or more complications during an inpatient hospital stay. The network graph highlighted renal failure without dialysis and septicemia and severe infections as highly connected complications in this population.

## Background

Since the inception of the Triple Aim, healthcare has endeavored to measure quality of care (Institute of Medicine, 2000). Complications during inpatient acute care are one aspect of quality of care (Leape, 1994). Acute care hospitals are faced with numerous types of complications including hospital-associated infections, medication errors, and falls (Centers for Medicare & Medicaid Services, 2016). The plethora of complications to be addressed by hospitals poses an operational challenge in the face of competing priorities.

The evolving field of data visualization continues to offer new approaches to understanding data. Network graphing analysis has been applied to numerous data sets (Viegas & Donath, 2004). Network graphing techniques allow a better way to understand and appreciate complicated relationships (Dekker & Colbert, 2004).

Network graphs can provide a quantitative framework for identifying complications with significant volumes and strong relationships to other complications, as an indicator of severity, as a method for prioritization of quality improvement work. Here we examine the application of network graphing techniques to acute care inpatient complications on acute care medical-surgical units of a quaternary care center.

## Methods

Data was retrospectively analyzed for patients discharged from medical-surgical units at Abbott Northwestern Hospital (Minneapolis, Minnesota) between January and October 2015. Patient level data from the electronic medical record is stored in an electronic data warehouse (EDW). We applied 3M Potentially Preventable Complications software (PCC) (3M, St. Paul, Minnesota) to this data to identify complications and accessed the results through a QlikView dashboard (Qlik, Radnor, Pennsylvania)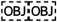. Patients who had two or more complications during a single inpatient stay were used to develop nodes and edge weights. We used Gephi (Paris, France) software to build the network graph (Gephi, 2016)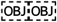.

Each node represents an individual complication and is color coded by PPC complication group. Each edge, or line, represents at least one patient that had both complications; thicker edges represent a more patients with both complications. The arrowheads are an artifact of the software and should be disregarded. In figure 1 each node is labeled with the number corresponding to the PPC description in table 1.

**Figure 1.**
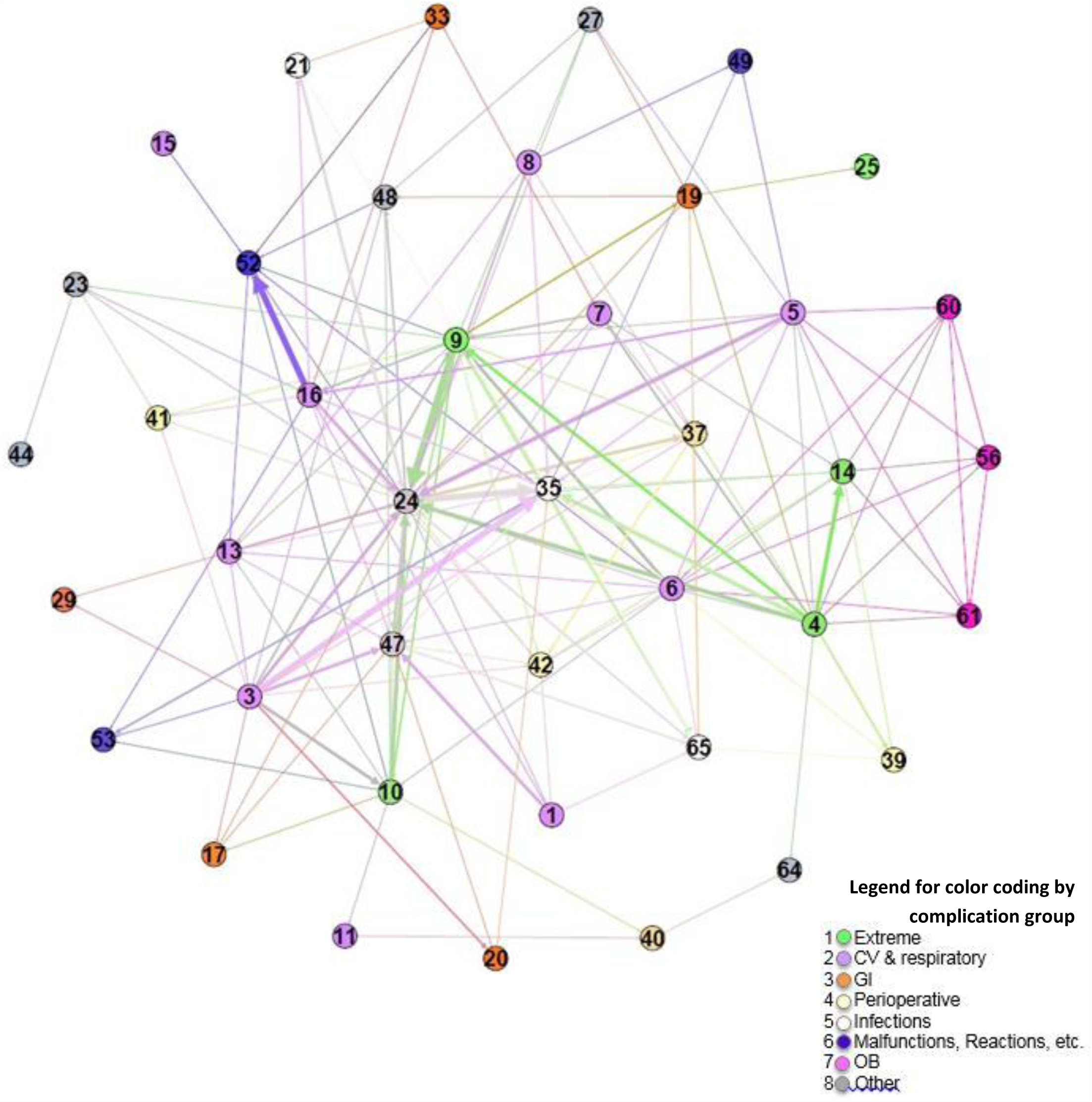
Network graph of complications in medical surgical patients.

**Table 1.** Complications

| **3M PPC Category** | **Number of patients w/ PPC** | **Edges** |
| --- | --- | --- |
| 1: Stroke & Intracranial Hemorrhage | 3 | 7 |
| 3: Acute Pulmonary Edema and Respiratory Failure without Ventilation | 18 | 27 |
| 4: Acute Pulmonary Edema and Respiratory Failure with Ventilation | 14 | 27 |
| 5: Pneumonia & Other Lung Infections | 9 | 17 |
| 6: Aspiration Pneumonia | 4 | 16 |
| 7: Pulmonary Embolism | 8 | 9 |
| 8: Other Pulmonary Complications | 3 | 7 |
| 9: Shock | 19 | 38 |
| 10: Congestive Heart Failure | 8 | 18 |
| 11: Acute Myocardial Infarction | 1 | 2 |
| 13: Other Cardiac Complications | 4 | 10 |
| 14: Ventricular Fibrillation/Cardiac Arrest | 6 | 14 |
| 15: Peripheral Vascular Complications Except Venous Thrombosis | 1 | 1 |
| 16: Venous Thrombosis | 13 | 24 |
| 17: Major Gastrointestinal Complications without Transfusion or Significant Bleeding | 2 | 4 |
| 19: Major Liver Complications | 4 | 8 |
| 20: Other Gastrointestinal Complications without Transfusion or Significant Bleeding | 2 | 4 |
| 21: Clostridium Difficile Colitis | 7 | 8 |
| 23: GU Complications Except UTI | 2 | 5 |
| 24: Renal Failure without Dialysis | 34 | 60 |
| 25: Renal Failure with Dialysis | 1 | 1 |
| 27: Post-Hemorrhagic & Other Acute Anemia with Transfusion | 3 | 5 |
| 29: Poisonings except from anesthesia | 2 | 2 |
| 33: Cellulitis | 3 | 4 |
| 35: Septicemia & Severe Infections | 25 | 38 |
| 37: Post-Operative Infection & Deep Wound Disruption Without Procedure | 7 | 12 |
| 39: Reopening Surgical Site | 4 | 6 |
| 40: Post-Operative Hemorrhage & Hematoma without Hemorrhage Control Procedure or I&D Proc | 2 | 3 |
| 41: Post-Operative Hemorrhage & Hematoma with Hemorrhage Control Procedure or I&D Proc | 2 | 5 |
| 42: Accidental Puncture/Laceration During Invasive Procedure | 6 | 10 |
| 44: Other Surgical Complication - Mod | 1 | 1 |
| 47: Encephalopathy | 12 | 22 |
| 48: Other Complications of Medical Care | 5 | 7 |
| 49: Iatrogenic Pneumothrax | 2 | 3 |
| 52: Inflammation & Other Complications of Devices, Implants or Grafts Except Vascular Infection | 10 | 14 |
| 53: Infection, Inflammation & Clotting Complications of Peripheral Vascular Catheters & Infusions | 3 | 5 |
| 56: Obstetrical Hemorrhage with Transfusion | 1 | 6 |
| 64: Other In-Hospital Adverse Events | 2 | 2 |
| 65: Urinary Tract Infection | 8 | 14 |

## Data

106 unique patients were found to have two or more complications. The 3M PPC software identified 66 complications, 39 of which occurred in pairs. As illustrated in Table 1, each complication involves a varying number of patients with differing partner complications. There were no pairs of complications that always occurred together.

## Results

The edge between complication 16: Venous Thrombosis and complication 52: Inflammation & Other Complications of Devices, Implants or Grafts except Vascular Infection, is the strongest edge in the network with a strength of 17. Complication 25: renal failure without dialysis is highly related to other complications, with 60 edges among the 34 patients. Complication 35: Septicemia and Severe infections is also highly connected. The connectivity of complication 35: Septicemia and Severe infections supports the clinical knowledge; patients with sepsis are complex and at risk for additional complications (Society of Critical Care Medicine, 2016).

## Discussion

The inherent challenges of using administrative data in 3M PPC software analysis are well documented in quality improvement literature (Lawthers, 2000) (Geraci, 1999). Clinical experts offer their critiques of the 3M PPC software methodology itself and some have a strong basis (Hughes MD & al, 2006). To date, relevant literature has not coalesced around a single definitive source of definitions for any complication. The use of the 3M PPC software provides a consistent application of the methodology across cases.

## Conclusion

Use of network graphs to identify complications with high connectivity or centrality may provide a useful framework for identification and prioritization of high impact quality improvement efforts. Being able to target organizational quality and process improvement efforts on connected complications may reduce overall complications at a higher rate than efforts focused on improving complications with low connectivity.

